# Revisiting Aristotle’s observation on bees: High floral constancy is common among bees but it is shaped by the locally abundant flowering species

**DOI:** 10.1101/2024.10.28.614270

**Authors:** Saket Shrotri, Sukhraj Kaur, Viraj Nawge, S Sandhya, Rohan Dandavate, Vinita Gowda

**Author notes:** Corresponding authors Vinita Gowda, Tropical Ecology and Evolution (TrEE) Lab, Department of Biological Sciences, Indian Institute of Science Education and Research (IISER) Bhopal, Bhopal, Madhya Pradesh, India. 462066. Saket Shrotri, Tropical Ecology and Evolution (TrEE) Lab, Department of Biological Sciences, Indian Institute of Science Education and Research (IISER) Bhopal, Bhopal, Madhya Pradesh, India. 462066. Our novel observations are that bees display floral constancy to the most abundant plant species in their local foraging patches and not the most abundant species on the landscape.

## Abstract

Floral constancy is the tendency of a pollinator to sequentially visit flowers of the same species despite the availability of other rewarding plants. In a phytodiverse community, resource assurance may lead to pollinators displaying floral constancy to the most abundant plant species. We tested this by investigating if pollinators are floral constant on the abundant or the non-abundant plants within a seasonally flowering tropical community. We quantified floral constancy in three social (*Apis* spp.) and two solitary (non-*Apis*) Indian native bees using three approaches, that is by manually tracking the bees, analysing their pollen load, and examining pollen sacs of returning bees at their hive. We next examined in *Apis cerana indica* if constancy in individual bees translated to hive-level constancy. We found that in our community with distinct co-flowering patches, bees were constant to the most abundant species within a localised patch, and not to the most abundant species in the landscape. While the pollen loads from both the social and solitary bees suggested that they show high floral constancy (> 70% uni-dominant pollen), their values differed significantly (*p* < 0.0001). Finally, approximately 90% of individuals within a hive showed floral constancy (monolectic), but collectively, a hive displayed polylectic foraging. Our findings highlight that the foraging patterns of native pollinators has been understudied and is a critical first step towards connecting reproductive assurances to plant-pollinator dependencies in large landscapes.

## 1. INTRODUCTION

In “Τ□ν περ□ τ□ ζ□α □στορι□ν” i.e., *Historia Animalium* (Latin, 4th century BCE), Aristotle noted that “On each expedition, the bee does not fly from a flower of one kind to a flower of another, but flies from one violet, say, to another violet, and never meddles with another flower until it has got back to the hive.” (Aristotle 350 BCE; cited by Grant 1950). This phenomenon of the tendency of a pollinator to sequentially visit flowers of the same species during a single foraging trip, despite the availability of other rewarding plant species, is defined as floral constancy or pollinator constancy (Grant 1950; Free 1970; Bennett 1883; Waser 1986; Laverty 1994; Gegear and Thomson 2004; Flanagan et al. 2009; Ali et al. 2017; Bruninga-Socolar et al. 2022; Takagi and Ohashi 2024). Since this observation by Aristotle, in pollination ecology, it remains to be empirically tested if floral constancy is common across different bee species.

Our current understanding of floral constancy is largely based on studies from temperate habitats, focusing predominantly on the European honeybee, *A. mellifera* (Susic Martin and Farina 2016), and *Bombus* spp. (Heinrich 1976; Wilson and Stine 1996; Gegear and Laverty 2005). Given the rich diversity of bee taxa in tropical and subtropical landscapes, we quantified floral constancy in native bees and addressed the following questions: Is floral constancy common among native bees? Is it driven by the floral abundance of a species in a patch? Do individuals within a hive exhibit a similar degree of floral constancy?

Floral abundance is one of the plant traits that promote floral constancy in a pollinator. The role of floral abundance in driving floral constancy in wild habitats is predominantly known from *Bombus spp*. (Heinrich 1976; Shibata and Kudo 2020) and *A. mellifera* (Bennett 1883; Fowler et al. 2016). Ethological observations from these two species show that foraging patterns in these bees are abundance-driven, and associated with an improvement in their flower-handling time (Woodward and Laverty 1992; Sanderson 2006). This can be expected since search images are more likely to be formed for the most commonly encountered traits (Tinbergen 1960; Levin 1972; Heinrich 1975; Wilson and Stine 1996; Goulson 2000). Overall, floral abundance is known to have a positive effect on the visitation rates of different bee species, irrespective of the associated nectar traits or species identity of the plant species (Thomson 1981; Kunin and Iwasa 1996; Dauber et al. 2010; Fowler et al. 2016). Thus, quantifying floral abundance becomes a crucial metric to understand how it shapes floral constancy in any community.

In the Asian paleotropics, bees are recognised as the primary pollinators in most plant communities (Liow et al. 2001; Ballantyne et al. 2017). Although bees are considered generalists (Gegear and Laverty 2001), specialised foraging behaviour in the form of floral constancy has been reported in both social and solitary bees (Grant 1950; Free 1963; Waser 1986). In social bee colonies, individual bees, even members of the same colony are known to exhibit floral constancy towards different plant species, and this preference may change over time (Free 1963; Sharma 1970; Chaturvedi 1973; Heinrich 1976; 1979; Jhajj and Goyal 1979; Waser 1986). Such individual-level specialised foraging behaviour has been proposed to improve the diversity in the pollen diet of social bees, providing them with a nutritional advantage over solitary bees (Devkota et al. 2024). On the other hand, solitary bees tend to collect mixed nectar or pollen resources in a single trip despite the higher energetic costs associated with this behaviour (Williams and Tepedino 2003; Eckhardt et al. 2014). Considering these contrasting foraging patterns between social and solitary bees, it is crucial to understand whether floral constancy is a common trait across different bee species, as this has direct implications for pollen movement within a community.

Floral constancy is often critically examined in the context of pollen carried by the pollinators and their subsequent movement among plants. When a pollinator switches between flowers of different species (also known as floral inconstancy) during its foraging trip, it increases the proportion of heterospecific pollen transfer (HPT). Studies among congeners have shown that HPT can reduce conspecific pollen fertilisation due to stigma clogging, pollen loss to foreign flowers, and other physiological interferences, and consequently, affect the reproductive success of plants (Waser 1986; Ashman and Arceo-Gomez 2013; Tur et al. 2016; Malecore et al. 2021). Since floral constancy can significantly reduce HPT (Morales and Traveset 2008; Arceo-Gómez and Ashman 2011; Moreira-Hernández and Muchhala 2019), estimating the constancy of different pollinators will also provide insights into how reproductive assurance may be maintained within a landscape (Feinsinger et al. 1988; Morales and Travest 2008; Arceo-Gómez and Ashman 2011).

Free (1970) noted that “The constancy of a bee to one plant species can be judged either by observing it directly or by analysing its pollen loads.” To the best of our knowledge, floral constancy is commonly measured by the following three methods: i) by manually observing the sequential visits of a pollinator and using a transition matrix representing the number of transitions a bee makes from one species to another during its one foraging trip (i.e., Constancy Index or CI; Straw 1972; Heinrich 1975; 1979; Bennett 1883; Waser 1986; De Jagar et al. 2011; van der Niet et al. 2020; Bruninga-Socolar, et al. 2022); ii) by quantifying the total amount of pollen present on the pollinator’s body (i.e., pollen load; Pangestika et al. 2017; Somanathan et al. 2020), and iii) by quantifying pollen from pollen sacs and assessing percent pollen purity (Betts 1935; Free 1970; Bennett 1883; Martínez-Bauer et al. 2021). All three methods are tedious yet simple and involve either direct manual observation of sequential visits of a pollinator or where the pollinator’s visitation pattern is inferred from pollen collected from its body (indirect methods). In order to compare and contrast the presence of floral constancy among the social and solitary bees in our study, we used all the above three metrics.

## 2. MATERIALS AND METHODS

### 2.1 Study site and system

This study was conducted on the Kaas plateau (कास पठार, GPS co-ordinates of Kaas plateau, elevation 1225 m, area 3.37 sq. km; Fig. 1), a UNESCO natural world heritage site located in the Western Ghats, Maharashtra, India. This plateau is an elevated outcrop of red laterite rock with a thin soil layer that hosts seasonal, herbaceous plants (Watve, 2013). We studied five native bees found on the Kaas plateau from August to October 2021, which is the flowering season on the plateau and it coincides with the monsoonal rains in peninsular India.

**Fig. 1:**
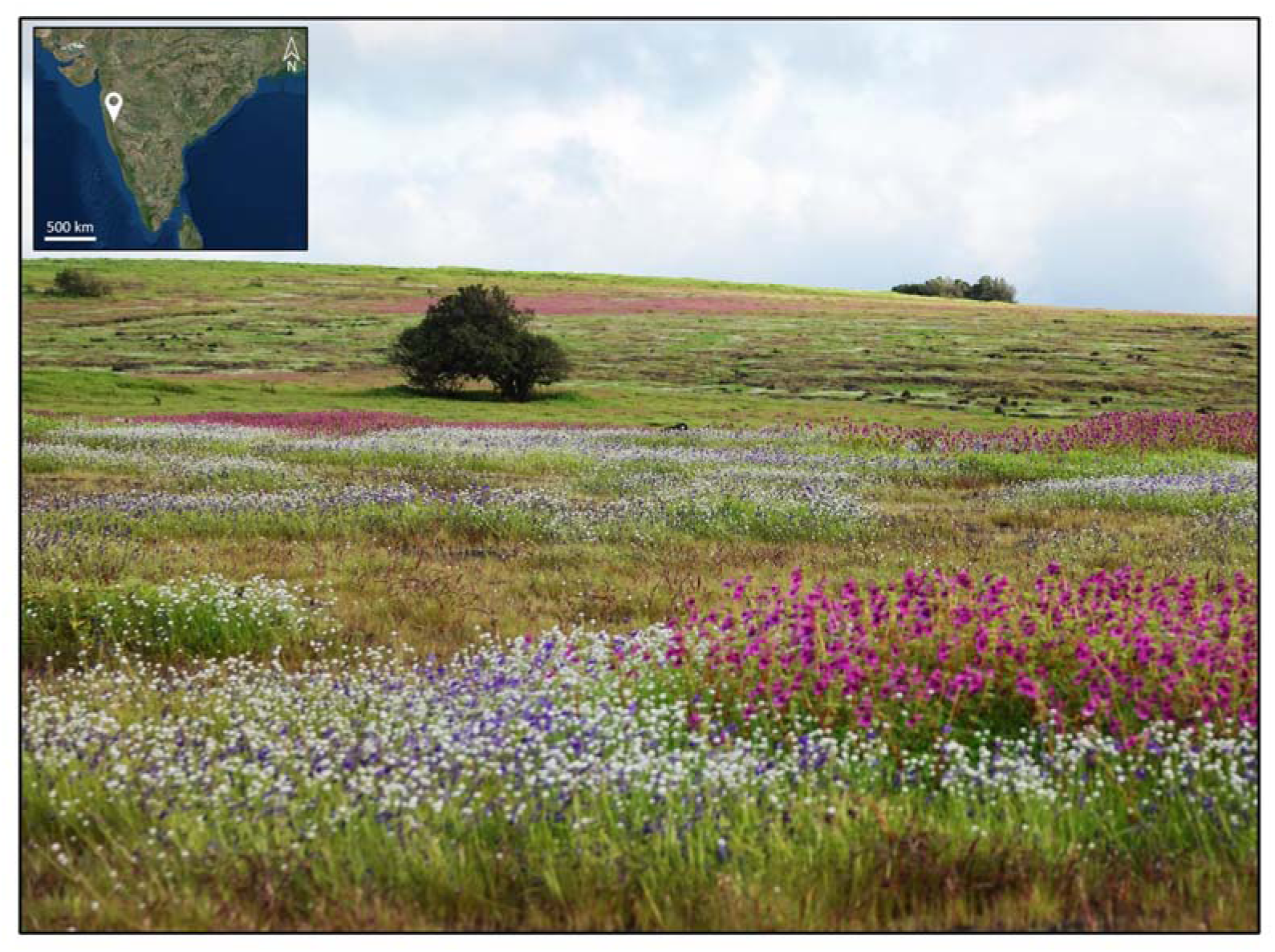
A landscape view of the seasonal herbaceous community found at Kaas plateau, Maharashtra. The inset map shows the relative location of Kaas plateau within Peninsular India.

The five native species investigated for their foraging behaviours in the wild are: three types of native honeybees (henceforth referred to as *Apis* bees) - the dwarf honeybee *A. florea*, the Indian honeybee *A. cerana indica*, and the giant honeybee *A. dorsata*, and two species of solitary bees (henceforth referred to as non-*Apis* bees) which are the blue-banded bee *Amegilla* sp. and an unidentified species, designated here as the ‘Small Black Bee’ species (SBB). We also carried out hive-level observations on the Indian honeybee *A. cerana indica* using bee boxes that were procured from Under The Mango Tree Society (UTMT) and installed at the study site in February 2021.

### 2.2 Pollen library preparation

Since the Kaas plateau plateau is known to have ∼ 180 species of angiosperms from ∼ 50 families (Fig. 1), to identify the pollen movement by bees, we first constructed a pollen library for all the plant species (n = 110) in the study site. For this, we collected pollen samples from unopened or recently opened flowers in absolute alcohol. Pollen grains were separated from the anthers through a process of agitation and centrifugation (Kearns and Inouye 1993) and imaged at 400X magnification using a light microscope (Leica DM2500, Leica, Germany), and using a Field Emission Scanning Electron Microscope (HR FESEM; ULTRA Plus, Zeiss with GeminiÒ column, Zeiss, Germany) available at IISER Bhopal.

### 2.3 Floral constancy of foraging trips and floral abundance

To standardise the measurement of floral constancy of pollinators we used a transition matrix representing foraging switches of a bee between any two plant species A and B (Online Resource Fig. S1; Bateman 1951). We next measured the Constancy Index (CI) for an individual bee as: Total number of intra-specific transitions ÷ Total number of observed transitions in its single foraging trip. Given this equation, the CI value ranged from zero (no constancy and no preference) to one (complete constancy and preference), with 0.5 representing indeterminable constancy. For each bee, we defined its “foraging trip” as a sequence of uninterrupted visits to several flowers where the pollen was not carried from one foraging trip to the next (Straw 1972).

Since documenting the entire foraging trip was logistically impossible on the plateau, transitions during foraging were measured for up to 25-30 consecutive visits within an observation plot for all the bees used in this study. In this way, we measured the CI values for a total of 1031 bees from *A. cerana indica* (n = 917), *A. dorsata* (n *=* 98), and *A. florea* (n *=* 16) separately, and the CI of a species was assigned by averaging the CI across all its individuals. All CI values were measured across 43 observation plots of size 4 to 5 sq. m from 08:00 am to 4:00 pm. The observation plot sizes were determined by the plant size and abundance and the ability to monitor pollinators for their consecutive visits during the same foraging trip. These plots were dispersed across the study site to capture the heterogeneous distribution of species and the unique combinations they formed. The Kaas plateau is known for its synchronous flowering phenology, with a few species exhibiting ‘mass flowering’ displays (Fig. 1; Anonymous et al., unpublished data). Therefore, while 33 plots contained at least 4 to 8 co-flowering species, 10 plots that contained mass flowering taxa had only two species in them. Next, we quantified floral abundance as the total number of flowers per plant species per plot. To eliminate the impact of interspecific variations in floral abundance across observation plots, we ranked each plant species within a plot based on their relative abundance on a scale of 1 to 8. Rank 1 meant the species had the highest floral abundance within a plot.

We used the Generalised Linear Mixed Models (GLMM), fitted by maximum likelihood with Poisson distribution, to test the effect of both floral abundance (fixed effect) and pollinator species (fixed effect) on the total number of bees with high floral constancy (CI values > 0.9). The identity of observation plots was the random effect in the GLMM. We used the *lme4* package (Bates et al. 2015) in R (version 4.3.0; R core team 2021) and the model was: Total number of bees ∼ Floral abundance rank + Bee species + (1| Observation plot).

### 2.4 Floral constancy via pollen load

We measured pollen load as the total amount of pollen carried on the body of a pollinator, which includes the pollen in the pollen baskets of the bee (Pangestika et al. 2017). For this, we attempted to collect a total of 50 foraging bees in absolute ethanol, to be later examined under a light microscope (Leica DM2500, Leica, Germany). Each pollen load was examined to identify pollen grains to their highest taxonomic level, with the help of reference images from the pollen library. Floral constancy for each bee was measured using separate metrics for pollen diversity and pollen abundance in each pollen load. Pollen diversity was measured as the number of plant species present in the pollen load. The total pollen abundance was measured as the sum of all pollen grains across all plant species per pollen load using a haemocytometer (Neubauer improved, Marienfeld Germany).

We grouped each pollen load based on their observed pollen diversity and abundance as pure pollen load (PP_L_) if 100% of the pollen grains were from the same plant species, uni-dominant pollen loads (DP_L_) if > 70% of the pollen grains were from the same plant species, and mixed pollen loads MP_L_ if < 70% of the pollen grains were from the same plant species. PP_L_ is equivalent to absolute pollen purity (Betts 1935; Free 1970). We compared pollen diversity, total pollen abundance, and pollen abundance of DP_L_ loads, between *Apis* and non-*Apis* bees using Welch’s t-test.

### 2.5 Floral constancy of individuals within a colony

To understand how the floral constancy of individual bees translates at the colony level, we quantified the floral constancy of worker bees within a colony. We established two colonies of *A. cerana indica* at two separate locations ∼ 5 km apart on the plateau. We used pollen traps to collect one pollen sac or corbiculae per individual from the tibia of returning worker bees. We collected ∼ 30 pollen sacs per bee box in absolute ethanol at a given time point, and this sampling was done for both boxes at 10-day intervals during September and October 2021. Pollen purity (PP_L_, DP_L_, and MP_L_) of each pollen sac was used as a proxy for the bee’s floral constancy and we used Pearson’s chi-square analysis to test the null hypothesis that ‘the number of pollen sacs is equally distributed among the three types of pollen purities.’ All plots and statistical analyses were performed in R (version 4.3.0; R core team 2021).

## 3. RESULTS

### 3.1 Bees displayed high floral constancy towards locally abundant species

The CI values for all 1031 observed bees ranged from 0.9 to 1.00, with the majority of bees (1021 bees or 99.03 %) demonstrating an absolute floral constancy (CI = 1, Table 1). Among the three *Apis* bees, all individuals of *A. dorsata* (100%) exhibited absolute constancy (CI = 1.0), followed by *A. cerana indica* (99.23%) and *A. florea* (81.25%; Table 1). The GLMM analysis revealed a positive correlation between a higher number of bees exhibiting floral constancy to plant species with higher abundance rank within the observation plots (z = 11.544, *p* < 0.001; Fig. 2; Online Resource Table S1). Thus, as the abundance of a plant species increased within an observational plot, there was a significant increase in the number of bees that were constant on that species. We found no effect of the taxonomic identity of bees on the relationship between floral abundance and the number of constant bees (Online Resource Table S1).

**Fig. 2:**
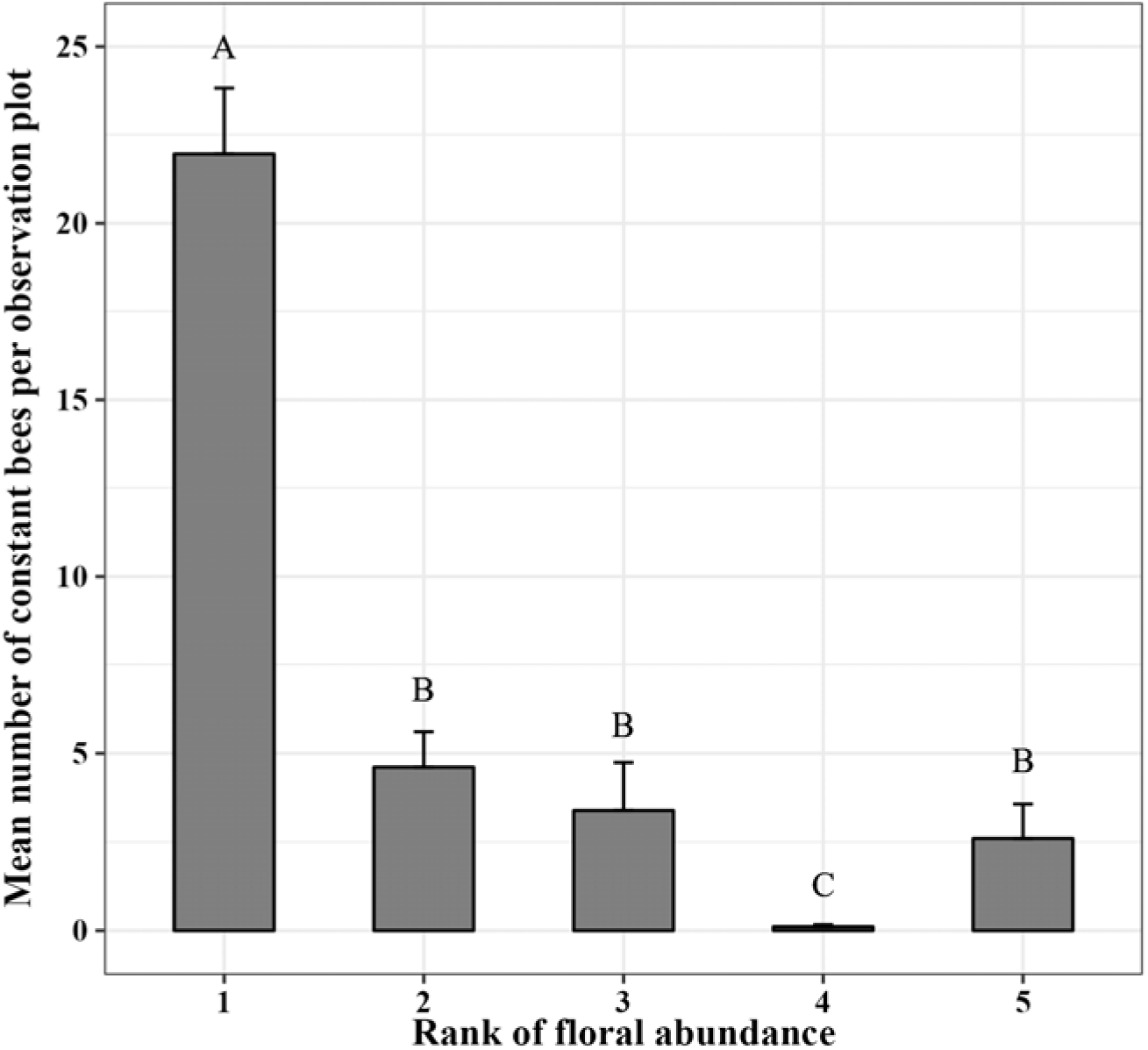
The number of bees (*n* = 1031) that were constant on a given rank of flowering species per observation plot (*n* = 43). No bees were observed on plants having rank abundance value > 5, that is lower floral abundances within the observation plot. The significant differences (*p* < 0.05) between ranks are depicted by alphabets on the respective bars. The statistical values are provided in Online Resource table S1.

**Table 1:** Floral constancy displayed by the three *Apis* bees across all observation plots (Constancy Index range: 0.9 to 1.0). Percent individuals along with the total individual counts per species is provided in parentheses. No bees were observed with a CI value < 0.9.

| Bee species<br>(Sample size) | % (individuals) displaying floral constancy |  |  |
| --- | --- | --- | --- |
|  | Constancy Index (CI) |  |  |
|  | 0.90 - 0.95 | 0.95 - 1.0 | 1.0 |
| *A. florea* (16) | 12.5% (2) | 6.25% (1) | 81.25% (13) |
| *A. cerana* (917) | 0.12% (2) | 0.55% (5) | 99.23% (910) |
| *A. dorsata* (98) | 0% | 0% | 100% (98) |
| Total (1031) | 0.29% (4) | 0.58% (6) | 99.03% (1021) |

### 3.2 *Apis* bees display higher floral constancy than non-*Apis* bees

We collected a total of 43 bees for pollen load measurements, out of which 20 were *Apis* bees (13 *A. cerana indica,* three *A. dorsata,* and four *A. florea*), and the remaining 23 were non-*Apis* bees (seven *Amegilla* sp. and 16 SBB; Fig. 3A). We found that only two individuals of *Apis* bees had pure pollen loads (PP_L_), while 41 out of 43 of the bees carried pollen from more than one plant taxa (ranging from 2 to 7 taxa per bee). Among the 41 bees carrying pollen from multiple taxa, 18 *Apis* bees and 13 non-*Apis* bees exhibited uni-dominant pollen loads (DP_L_). The remaining 10 bees displayed mixed pollen loads (MP_L_). We found no significant difference in the total pollen load (Welch’s t-test, df = 39.29, *p* = 0.5193) as well as in the pollen diversity (Welch’s t-test, df = -1.5248, df = 36.54, *p* = 0.1359) for *Apis* and non-*Apis* bees. However, the quantity of DP_L_ was significantly higher in the pollen loads of *Apis* bees compared to non-*Apis* bees (Welch’s t-test, df = 33.48, *p* < 0.0001).

**Fig. 3:**
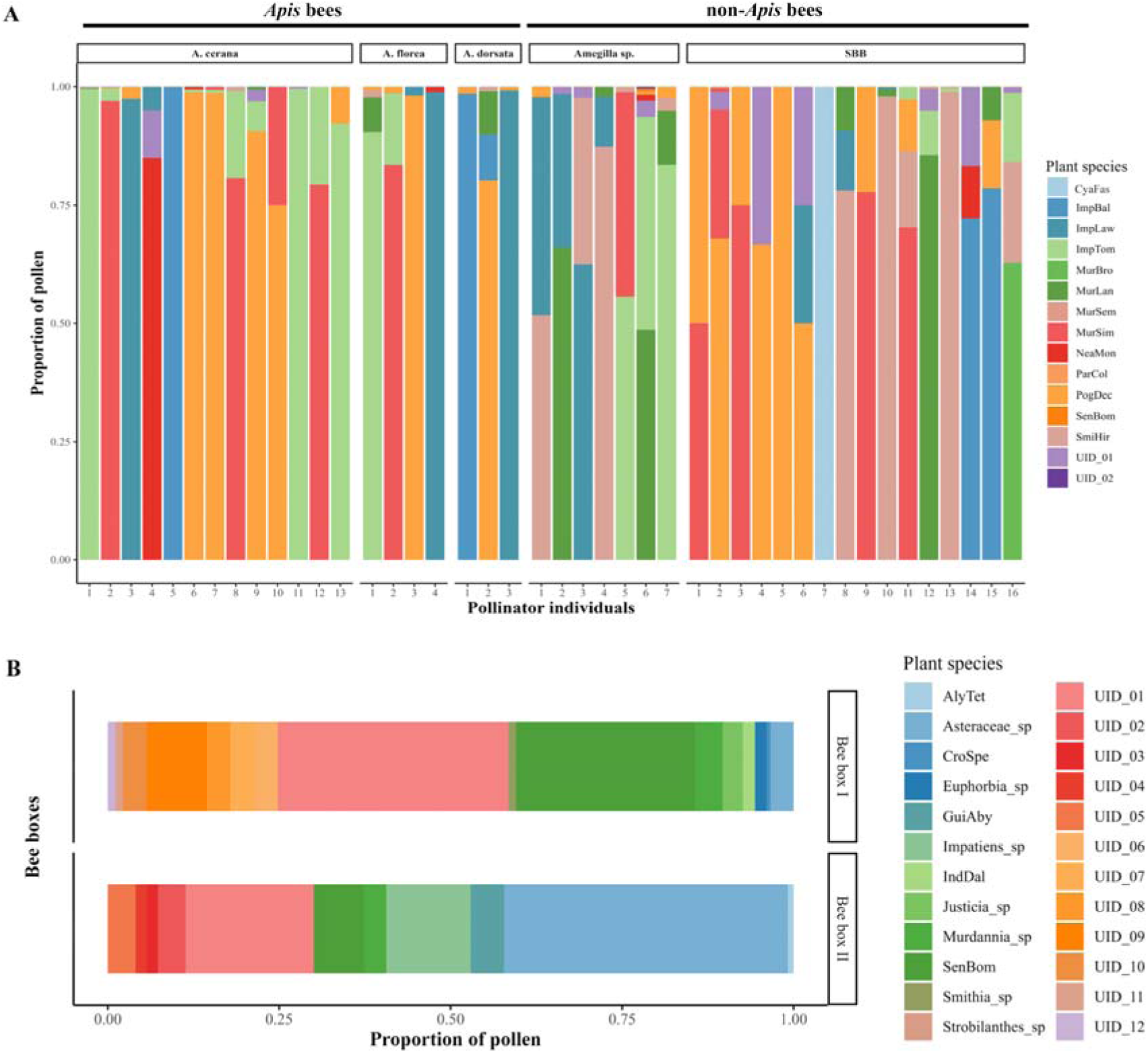
Floral constancy based on the pollen profile of bees. A) Pollen profile from pollen loads of *Apis* (20) and non-*Apis* (23) bees caught during foraging. B) Collective pollen profile from a total of 294 pollen sacs from two bee boxes of *Apis cerana indica*. For the pollen profile of individuals within a box and full plant species names, see Online Resource figure S2 and table S2, respectively.

### 3.3 Specialised individuals in generalised hives

We collected a total of 303 pollen sacs from two bee boxes at six different time points during the study. However, nine of these sacs were compromised during the process of sample collection, resulting in a final count of 294 samples (173 from bee box I and 121 from bee box II; Online Resource Fig. S2). During the first half of September, heavy rainfall limited our access to the field site, preventing the collection from bee box II during the first two time points.

We identified pollen from 24 different plant species in the pollen sacs collected from both bee boxes (18 from bee box I and 10 from bee box II; Fig. 3B). Interestingly, pollen from only four plant species was common in both boxes (Fig. 3B). Out of the 294 samples, 259 sacs (or 88.09%) were PP_L_ (i.e., one pollen type). The distribution of the PP_L_ pollen sacs in the two bee boxes ranged from 84.39% (146 pollen sacs) for bee box I to 93.38% (113 pollen sacs) for bee box II (Online Resource Fig. S2). The remaining 35 sacs (or 11.91 %) had DP_L_; they represented 15.61% (27 pollen sacs) in bee box I and 6.62% (eight pollen sacs; Online Resource Fig. S2) in bee box II. All DP_L_ samples had a maximum of only two types of pollen grains. No pollen sac was found to have MP_L_. Pearson’s chi-square test rejected the null hypothesis that pollen sacs within a hive show an equal distribution among the three pollen purity groups (χ2 = 4.6693, df = 1, *p* = 0.03071), indicating that bees are more likely to carry PP_L_ than DP_L_ or MP_L_.

## 4. DISCUSSION

In this study, we examined floral constancy among the five Indian native bees in a seasonal herbaceous community to understand the impact of foraging behaviour on pollen movement. We used three different methods to quantify floral constancy that was a) monitoring the visitation sequence of bees, b) assessing the purity of pollen loads carried by bees during foraging, and c) examining the pollen sacs post-foraging at the hives. We first calculated floral constancy using the constancy index (CI; Online Resource Fig. S1) that was modified from the Bateman’s index (BI; Waser 1986) because the Bateman’s index value is undefined when a pollinator visits only one species in a foraging trip and therefore shows null value in three out of the four transitions within the Bateman’s matrix (Gegear and Laverty 2005; Online Resource Fig. S1). Bateman identified this condition to represent an already determinable ‘preference’ by a pollinator and therefore did not quantify constancy. However, it has been shown that preferences can be labile among individuals within a species (Waser 1986), hence we used the CI metric to include individual-level variations in floral constancy. The pollen diversities found in the bee’s pollen load and pollen sacs (Fig. 3) were predictable from the floral constancy values calculated from the bees’ foraging trips using the CI metric (Online Resource Fig. S1; Table 1). Thus, our results also validate the reliability of the CI metric as a measure of floral constancy and its usefulness in predicting pollen removal by bees from only manual observations, and the use of pollen loads as a possible prediction for putative foraging pathways the pollinator may have taken.

### 4.1. Floral constancy is common among native bees

Our observations of visitation sequences of bees revealed a high degree of floral constancy (CI > 0.90; Table 1) across all *Apis* species (in this study: *A. florea*, *A. cerana indica*, and *A. dorsata*) suggesting very rare switches by a bee during its foraging trip. Since our observation plots were spread across the plateau and were variable in their plant diversities, this also suggests that the bees display similar floral constancy regardless of the distribution of plant species across the plateau and are polylectic (Fig. 3). Similar high floral constancy values among social bees such as *Apis* and *Bombus* have also been reported in *A. mellifera* (Bennett 1883; Susic Martin and Farina 2016), *B. vagans* (Heinrich 1976; 1979; Wilson and Stine 1996), and *B. impatien*s (Gegear and Laverty 2005). Floral constancy models predict that the pollen removal or pollen diversity in the pollen sac will be biassed towards the species on which pollinators show high floral constancy (Bennett 1883; Betts 1935; Martínez-Bauer et al. 2021). We noted this bias in individuals of *A. cerana indica* that were returning to their hives, where ∼ 90% of the pollen sacs from individual bees consisted of pollen from a single plant species. The plant species found in the hives were also observed to be the ones that were visited by bees in our manual observations and that were present in the pollen load counts (CI > 0.9; PP_L_ or DP_L_; Fig. 3B; Online Resource Fig. S2). The high proportion of individuals with pure pollen loads also means that heterospecific pollen was low (Fig. 2A; Online Resource Fig. S2) and this is also indicative of specialisation by bees within a hive towards a few plant species (Smith et al. 2019; Online Resource Fig. S2). Some of these plant species were *Impatiens* sp., *Justicia diffusa*, *Senecio bombayensis*, and *Murdannia simplex*. These plant species were also found in the pollen loads of both *Apis* and non-*Apis* bees in high proportions (∼ 77% pollen loads with DP_L_; Fig. 3A). To the best of our knowledge, our results are the first to show that the high pollen purity and low heterospecific pollen found in the pollen sacs and pollen loads of the three species of native *Apis* bees can be explained by the high floral constancy they display in their foraging trips (high CI values).

### 4.2. Bees are constant on the locally abundant flowering species

Floral constancy is an adaptive behaviour and is known to be influenced by the local abundance of floral resources (Heinrich 1979; Kunin 1993; Dauber et al. 2010; Crone 2013; Pangestika et al. 2017). Previous studies have recorded floral constancy in *A. cerana indica*, from managed habitats (Chaturvedi 1973; Jhajj and Goyal 1979; Suryanarayan et al. 1992). However, these studies primarily focused on plantations dominated by a single plant species, and hence a single pollen type may have been recovered. Moreover, since floral constancy in these studies was assessed solely through pollen loads and not supported by manual observations of sequential visits, they do not provide insights into the foraging behaviour of *A. cerana indica* in a wild habitat with heterogeneous distribution of plant species.

Results from our manual observations reveal that *Apis* bees tend to be constant on the most abundant flowering species within a localised foraging patch which refers to the observation plots in our study (Fig. 2; Online Resource Table S1). This means that while different individuals of the same species visited different flowers, a higher proportion of bees were constant on the locally abundant species within a foraging patch (plant species with rank abundance = 1). Floral constancy as a strategy is supposed to have evolved due to the formation of search images based on the commonly encountered traits which enhance the efficiency of flower detection, and reduce the flower handling and travel time between flowers, thereby minimising the energetic costs associated with foraging (Lewis 1986; Woodward and Laverty 1992; Kunin and Iwasa 1996; Wilson and Stine 1996; Goulson 2000). Our observed constancy in *Apis* bees to the locally abundant species suggests that the native bees may be optimising reward gain by foraging on the most common species within a patch.

Although several mass flowering species were present on the Kaas plateau, the observation plots in this study were distributed such that 33 of the 43 plots did not contain any mass flowering species. The presence of floral constancy towards the locally abundant plant species suggests that reproductive benefits from floral constancy are not limited to a few mass flowering species but are also extended to the locally abundant but rare taxa. Thus, the relative abundance rather than the absolute abundance of a species was critical in determining the floral constancy of a pollinator within a patch. This has been shown in bumblebees (*B. ignitus*) where floral constancy increased with patchy distributions of flowers (Takagi and Ohashi 2024). Theoretical models have also shown that floral constancy can be expected towards rare species as long as they maintain a certain threshold density (Kunin and Iwasa 1996; Hayes & Grüter 2023). We propose that in seasonally flowering herbaceous communities such as in the Kaas plateau, rare species may be ensuring reproductive fitness and avoiding pollen limitation via improved pollinator visitations through floral constancy by creating locally abundant floral displays.

### 4.3. *Apis* bees show higher floral constancy than non-*Apis* bees

Foraging behaviours can maintain pollen diversity within a hive and are known to be affected by nutritional requirements (Linsley and MacSwain 1958; Vaudo et al. 2016; 2020) and life histories of the social and solitary bees. For instance, individuals within a social colony will achieve nutritional requirements by collective foraging through recruitment mechanisms (Seeley 1989; Frisch 1993), whereas solitary bees rely on an individual’s foraging experiences (Heinrich 1976; Williams and Tepedino 2003; Eckhardt et al. 2014). Consequently, solitary bees will exhibit more interspecific transitions than social bees (Ne’eman et al. 2006; Biddinger et al. 2013; Smith et al. 2019). In our study, all *Apis* bees were social bees and abundant in the community, while all the non-*Apis* bees were solitary and relatively less abundant. Analysis of pollen loads from both groups revealed a higher prevalence of uni-dominant pollen in social bees compared to solitary bees (Fig. 3A). This distinction can be attributed to the differences in the nutritional requirements of the two groups (Vaudo et al. 2016; 2020) and suggests higher interspecific transitions among the non-*Apis* bees. Pollen loads are indicative of foraging behaviour and floral constancy, both of which are known to be adaptive in bees (Free 1970, Ne’eman et al. 2006; Grüter & Ratnieks 2011). We propose that the presence of uni-dominant pollen in both social and solitary bees is suggestive of floral constancy having an adaptive value in this hyperdiverse, seasonal community.

### 4.4. Specialist individuals make generalist colonies

Despite the rich diversity of bee taxa in tropical and sub-tropical landscapes, our knowledge of the foraging behaviours of individual bees within a hive is still limited and mostly known from *B. terrestris* (Spaethe & Weidenmüller 2002) and *A. mellifera* (Free 1963; Heinrich 1976). We show that in *A. cerana indica* hives ∼ 90% of the individuals carry uni-dominant pollen and heterospecific pollen diversity is very low, that is, they show high floral constancy towards one plant species. However, individuals from a hive collectively visited 4-10 species within a day (Online Resource Fig. S2). Thus *A. cerana indica* is polylectic as a species, its hive is a generalist colony with individuals showing labile preferences, but the individuals are specialised with high floral constancy towards a single plant species.

A floral visitor that remains constant during its foraging trip is likely to be an effective pollinator, as the foraging sequence shapes the pollen movement and reduces heterospecific pollen transfers (Baker and Hurd 1968; Ghazoul 2005; van der Niet et al. 2020). Therefore, a comprehensive understanding of the floral constancy of different bee species will eventually help identify their role in the reproductive assurances of plants within a hyperdiverse landscape where they forage. While temporal factors such as the lifespan of the forager and flowering phenology durations can influence foraging energetics and shape the floral constancy of a pollinator (Heinrich 1976; Bruninga-Socolar et al. 2022; Takagi and Ohashi 2024), we do not think that these will be critical in the seasonal communities such as the Kaas plateau.

Finally, we think Aristotle’s remark may have been inspired by his observations on a landscape similar to our study site and we agree that on each expedition nearly all the bee species we studied on Kaas plateau too did not fly from a flower of one kind to a flower of another, but instead stayed with its first chosen flower type, and never meddled with another flower until it got back to its hive, and thus maintained high floral constancy.

## ACKNOWLEDGMENTS

Authors thank the Ministry of Human Resource Development (MHRD) start-up grant to VG via IISER Bhopal. VG thanks BD for being an understanding, patient, and personal support throughout this study. SS1 (Saket Shrotri) thanks DST for the INSPIRE fellowship (IF190233) and INLAKS Ravi-shankaran foundation for INLAKS small grant 2020; SK thanks Council of Scientific and Industrial Research for the Senior Research Fellowship (09/1020(0186)/2019-EMR-1). We thank IISER Bhopal for infrastructure, academic, and research support, especially SEM facility. We thank PCCF, the Maharashtra Forest Department, the DFO of Satara Division, and the Joint Forest Management Committee (JFMC), Kaas, Satara for research and collection permits. We thank the staff from UTMT society (Under The Mango Tree) for helping with bee box provision and training to SS, SK, and VG. We thank the local beekeepers Mr. R. K. Atale, Mr. Datta Kokre and Mr. Dattatray Kokre. We also thank Ms. Ritu Yadav for her assistance during the setup of the bee boxes.

## DECLARATIONS

## Funding

This study was primarily funded by small students grant for conservation by INLAKS Shivdasani Foundation. Rest of the funding was provided by Ministry of Human Resource and Development, Government of India.

## Conflict of interest

Authors have no conflict of interest to declare.

## Ethical approval

Not applicable

## Consent to participate

Not applicable

## Consent to publication

Not applicable

## Availability of data and material

The data will be deposited in Dryad data repository once the manuscript is accepted for publication, or upon request by the reviewers.

## Code availability

The codes used in this manuscripts will be deposited in Dryad data repository once the manuscript is accepted for publication, or upon request by the reviewers.

## Authors’ contribution

VG, SS1 (Saket Shrotri), SK, and VN conceptualised the study. VG and SS acquired the funds and administered the project. SS1, SK, VN, SS2 (S Sandhya), and RD carried out fieldwork and collected data. SS1 and SK carried out all formal analyses. VG, SS1, and SK wrote the original draft and edited drafts were shared with VN, SS2, and RD for comments. All authors approved the final version of the manuscript and agreed to be held accountable for the content therein.

## Notes

### Competing Interest Statement

The authors have declared no competing interest.

