## Supplementary files for "Revisiting Aristotle’s observation on bees: High floral constancy is common among bees but it is shaped by the locally abundant flowering species"

**Table S1:**

Fixed effects table for the GLMM fitted to test the effect of flowering abundance and bee species on number of constant bees. The significance level is set at p < 0.05.

|  |  | **Estimate** | **SE** | **z** | ***p*** |
| --- | --- | --- | --- | --- | --- |
|  | Intercept | 2.9267 | 0.2532 | 11.544 | < 0.001 |
| Rank | Rank 2 | -1.5601 | 0.0851 | -18.333 | < 0.001 |
|  | Rank 3 | -1.6869 | 0.1369 | -12.326 | < 0.001 |
|  | Rank 4 | -4.8880 | 0.9989 | -4.893 | < 0.001 |
|  | Rank 5 | -1.3333 | 0.2998 | -4.448 | < 0.001 |
| Bee species | *Apis dorsata* | -0.1034 | 0.1235 | -0.836 | 0.403 |
|  | *Apis florea* | 0.0264 | 0.3005 | 0.088 | 0.930 |

**Table S2:**

Taxonomic names for six letter species codes of flowering species present in: A) Pollen load data, and B) Pollen sac data. The flowering status of the respective plant species in terms of their flowering abundance is represented as MF (mass flowering), or nMF (non-mass flowering).

|  | **Sr. No.** | **Flowering species** | **Abbreviation** | **Flowering status** |
| --- | --- | --- | --- | --- |
| A | 1 | *Cyanotis fasciculata* | CyaFas | MF |
|  | 2 | *Impatiens balsamina* | ImpBal | nMF |
|  | 3 | *Impatiens lawii* | ImpLaw | nMF |
|  | 4 | *Impatiens tomentosa* | ImpTom | nMF |
|  | 5 | *Murdannia crocea* | MurCro | nMF |
|  | 6 | *Murdannia lanuginosa* | MurLan | nMF |
|  | 7 | *Murdannia semiteres* | MurSem | MF |
|  | 8 | *Murdannia simplex* | MurSim | nMF |
|  | 9 | *Neanotis montholonii* | NeaMon | nMF |
|  | 10 | *Paracaryopsis coelestina* | ParCoe | nMF |
|  | 11 | *Pogostemon deccanensis* | PogDec | nMF |
|  | 12 | *Senecio bombayensis* | SenBom | nMF |
|  | 13 | *Smithia hirsuta* | SmiHir | nMF |
|  | 14 | Unidentified | UID_X | - |
| B | 1 | *Indigofera dalzellii* | IndDal | nMF |
|  | 2 | *Senecio bombayensis* | SenBom | nMF |
|  | 3 | *Justicia diffusa* | JusDif | nMF |
|  | 4 | *Murdannia* sp. | Mur_sp | nMF |
|  | 5 | Asteraceae (excluding SenBom) | Asteraceae | nMF |
|  | 6 | *Euphorbia* sp. | Eup_sp | nMF |
|  | 7 | *Smithia* sp. | Smi_sp | nMF |
|  | 8 | *Crotolaria spectabilis* | CroSpe | nMF |
|  | 9 | *Strobilanthes* sp. | Str_sp | nMF |
|  | 10 | *Guizotia abyssinica* | GuiAby | nMF |
|  | 11 | *Impatiens* sp. | Imp_sp | nMF |
|  | 12 | *Alysicarpus tetragonolobus* | AlyTet | nMF |

**Figure S1:**

Transition matrix given by Bateman (1951) and the formula used to calculate Constancy Index (CI) in our study. Numerals 1 and 2 refer to plant species being visited by a pollinator, and alphabets A to D refer to all types of possible visitation transitions between these two plant species. CI was calculated for each foraging trip separately.

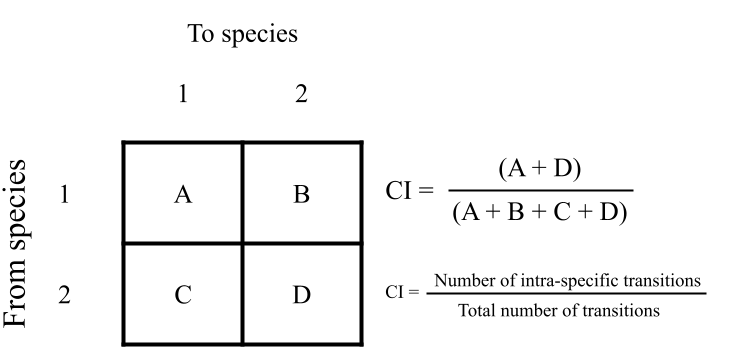

**Figure S2:**

Pollen profile of *Apis cerana indica* across bee boxes I and II. A) Individual-level pollen profile Time points for pollen sac collection from September (Sep) to October (Oct) are represented in different facets from left to right. For plant species codes, see Appendix S4. B) The distribution of pollen sacs with respect to their pollen purity (PP_L_ in grey and DP_L_ in black) for bee boxes I and II across different collection time points (September to October).

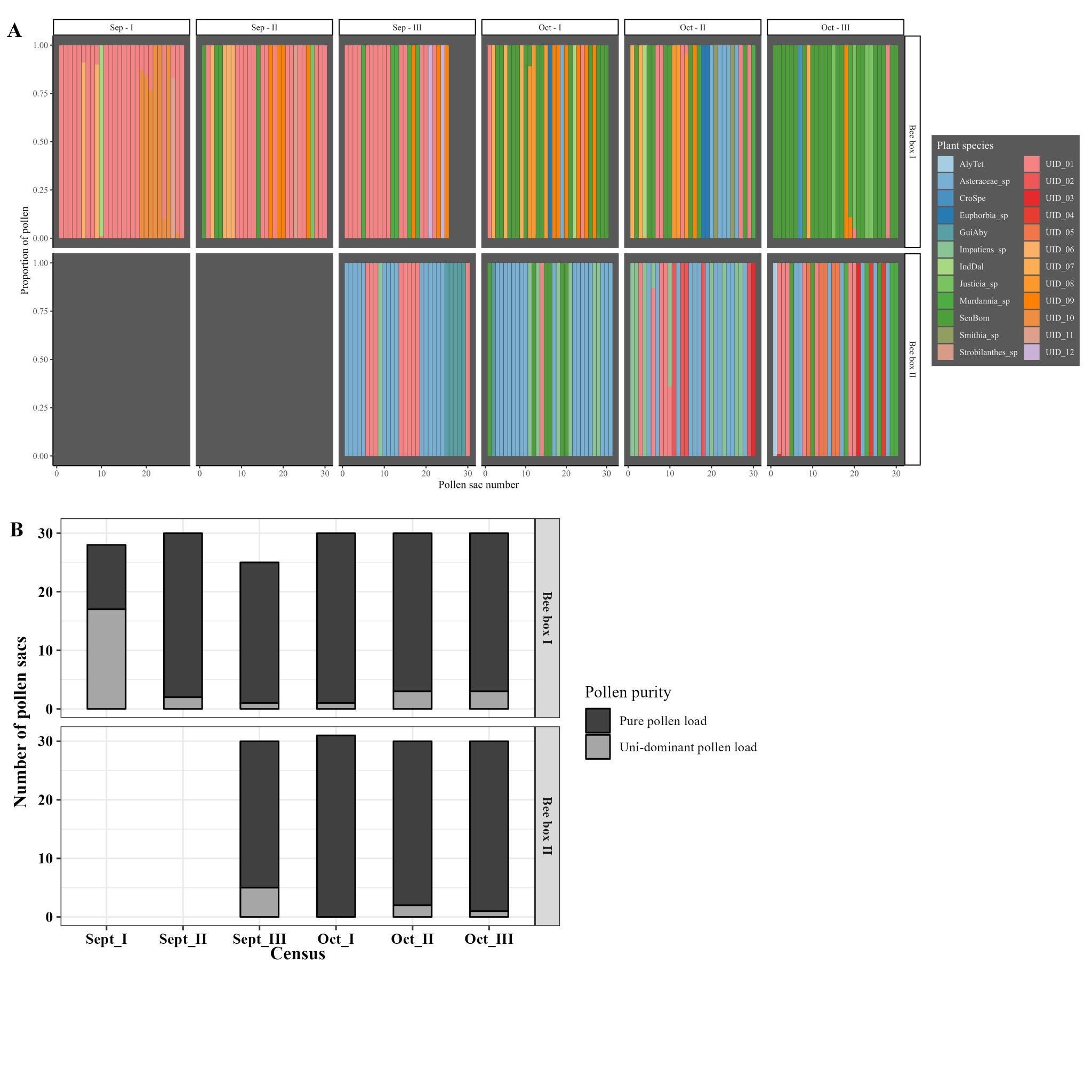
